# Epidermal γδ T cells, CD8 T cells and macrophages are increased in number in alopecia areata and express BST2 as part of an interferon-driven antiviral gene signature

**DOI:** 10.1101/2025.06.26.661822

**Authors:** Jake Coast, Alex Gonzalez, Cristina Velasquez, Milana Kansky, Yosiris Valdovinos-Hagan, Matilde Macedo, Andrea Solano, Antolette Kasler, Julie M. Jameson

**Affiliations:** Department of Biological Sciences, California State University San Marcos, CA 92096

## Abstract

Alopecia areata is an autoimmune disorder affecting approximately 2% of the global population, characterized by immune-mediated disruption of hair follicle immune privilege and unpredictable hair loss. While CD4 and CD8 T cells are established drivers of pathogenesis, the roles of other immune cell populations remain incompletely defined. Here, we reaffirm the pathogenic role of CD8 T cells and identify a contribution of epidermal γδ T cells and macrophages to alopecia areata. Using publicly available single-cell RNA sequencing (scRNAseq) data from the skin of C3H/HeJ mice with alopecia areata, we demonstrate that epidermal γδ T cells upregulate genes associated with immune homeostasis, proliferation, and inflammation, including *BST2*, an interferon-stimulated antiviral protein. In vivo, epidermal γδ T cells and keratinocytes are increased in number, with *BST2* expression enriched around hair follicles. BST2 is also expressed on CD8 T cells actively producing IFN-γ and on CD11b^+^ macrophages with signatures of antiviral and complement system pathways. In human alopecia areata skin, *BST2* expression is elevated in alopecia areata and downregulated following treatment with the JAK inhibitor tofacitinib. Together, these findings position BST2 as a marker of interferon-driven immune activation and highlight epidermal γδ T cells as a contributor to alopecia areata pathogenesis.

## Introduction

Alopecia areata is an autoimmune disorder characterized by spontaneous, patchy hair loss, affecting approximately 2% of the global population (Lee et al. 2020). The disease can progress to alopecia totalis, complete hair loss on the scalp, or alopecia universalis (near complete or complete body hair loss) (Simakou et al. 2019). T cells play a central pathogenic role in alopecia areata by disrupting immune privilege in the hair follicle (Connell and Jabbari 2022). In particular, CD8 T cells in both humans and mice with alopecia areata are sufficient to transfer disease in mouse models and human xenograft studies (Gilhar et al. 1998; McElwee et al. 2005; Xing et al. 2014). CD4^+^CD25^-^ T cells can transfer systemic disease, while CD4^+^CD25^+^ T regulatory cells can protect from disease onset (Han et al. 2015). CD4^+^CD25^-^ T cells can transfer systemic disease, while CD4^+^CD25^+^ T regulatory cells can protect from disease onset (McElwee et al. 2005). However, the contributions of other skin-resident populations such as γδ T cells, remain poorly understood.

Epidermal γδ T cells participate in wound repair, tumor surveillance, and maintenance of skin homeostasis (Johnson et al. 2020; Sharp et al. 2005; Toulon et al. 2009). Dermal γδ T cells also contribute to hair follicle neogenesis via FGF-9 production and to psoriasis pathogenesis through IL-17 secretion (Cai et al. 2011; Gay et al. 2013). In humans, Vδ1^+^ γδ T cells are found nears the hair follicle in patients with alopecia areata; where they exhibit higher expression of NKG2D and IFN-γ suggesting a pro-inflammatory phenotype (Uchida et al. 2020). γδ T cells interact with resident and infiltrating antigen presenting cells (APCs) including macrophages through cytokine and chemokine signaling (Cai et al. 2011; Nielsen et al. 2017). Macrophages also infiltrate the perifollicular and perivascular regions in patients with alopecia areata (Nasiri et al. 2020; Ranki et al. 1984). In both humans and mice with alopecia areata, CD4 and CD8 T cells exhibit clonal expansion and IFN-γ dominant gene signatures(Borcherding et al. 2022). Notably, macrophages exhibit an M1 polarization phenotype, which is induced by IFN-γ (Borcherding et al. 2022).

Genome-wide association studies further support an IFN-γ-driven mechanism in alopecia areata identifying susceptibility loci associated with T cells, including genes in the histocompatibility complex (MHC) and MHC Class I-like genes *UBLP3*/*ULBP6* and *MICA1/2,* cytokines such as *IL-2/IL-15/IL-21/IFN-*γ, and regulators of T cell activity such as *CTLA4/JAK/STAT* (Jagielska et al. 2012; John et al. 2011; Martinez-Mir et al. 2007; Petukhova et al. 2010). Small molecule inhibitors targeting IL-15 and IFN-γ downstream effectors, such as Janus kinases (JAKs), reverse hair loss in mouse models and patients with alopecia areata (Jabbari et al. 2016; Kwon et al. 2023; Xing et al. 2014). Furthermore, gene expression signatures in JAK inhibitor-treated patients stratify responders and nonresponders, offering potential for personalized treatments (Crispin et al. 2016).

In this study we investigated the role of γδ T cells and the immune cells they regulate in alopecia areata to uncover novel gene signatures and cellular interactions in the disease. We used publicly available single cell RNA sequencing (scRNAseq) data from C3H/HeJ mice to show that in alopecia areata, γδ T cells upregulate a set of genes that are distinct from αβ T cells.

Epidermal γδ T cells exhibit an antiviral gene signature, including bone marrow stromal antigen- 2 (BST2), an antiviral protein tetherin as well as pathways associated with cell proliferation (Tiwari et al. 2019). *In vivo* we observe an increased number of epidermal γδ T cells and keratinocytes in C3H/HeJ mice with alopecia areata. BST2 is also expressed by infiltrating CD8 T cells and CD11b^+^ macrophages in alopecia areata. Additionally, we found that *BST2* is downregulated in alopecia areata patients treated with JAK inhibitors. Together, these findings reveal a BST2-driven antiviral response in both skin-resident and infiltrating immune cell populations in alopecia areata skin.

## Results

### γδ T cells upregulate genes involved in immune cell homeostasis, proliferation, and inflammation in alopecia areata

γδ T cells reside in the epidermis, including the infundibulum region of the hair follicle, a key site impacted in alopecia areata (Uchida et al. 2020; Zhang et al. 2013). To delineate which genes are differentially expressed by epidermal γδ T cells in alopecia areata, we reanalyzed publicly available single-cell RNA sequencing data from C3H/HeJ mice with or without alopecia areata (Borcherding et al. 2022). An unsupervised clustering analysis of the 4,141 single cells using Loupe Browser identified nine distinct cell clusters. Each cell had an average of 3,323 median UMI counts per cell and 93% of the fractions reads for each cell were kept. Populations of clusters were manually annotated using the analysis of expression patterns of canonical markers for T cells (*Cd3d, Cd3g, Cd3e, Cd28, Cd8a, Cd4*), APCs (*Itgax, Itgam, Xcr1, Cd207*), B cells (*Cd79a, Cd79b, Cd19*), NK cells (*Klrc1, Klrc2, Nkg7*), macrophages (*Cd14, Lyz2*), keratinocytes (*Krt5, Krt14*), and fibroblasts (*Lum, Dcn*) (Fig. 1A). UMAP populations were split into control and alopecia areata groups to identify the relative percentage of each cell lineage for each condition. We clustered T cells into αβ T cells (*CD3ε, Trac*, *Trbc1*, *Trbc2*), γδ T cells (*CD3ε*, *Trdc*, *Tcrg-C1*, *Trdv4*), and Vγ5Vδ1 epidermal γδ T cells (*CD3ε, Trdv4*, *Tcrg-V5*) to characterize cellular subpopulations and gene expression (Fig. 1B). As previously reported, there was a marked increase in αβ T cells in mice with alopecia areata (Borcherding et al. 2022).

**Figure 1.**
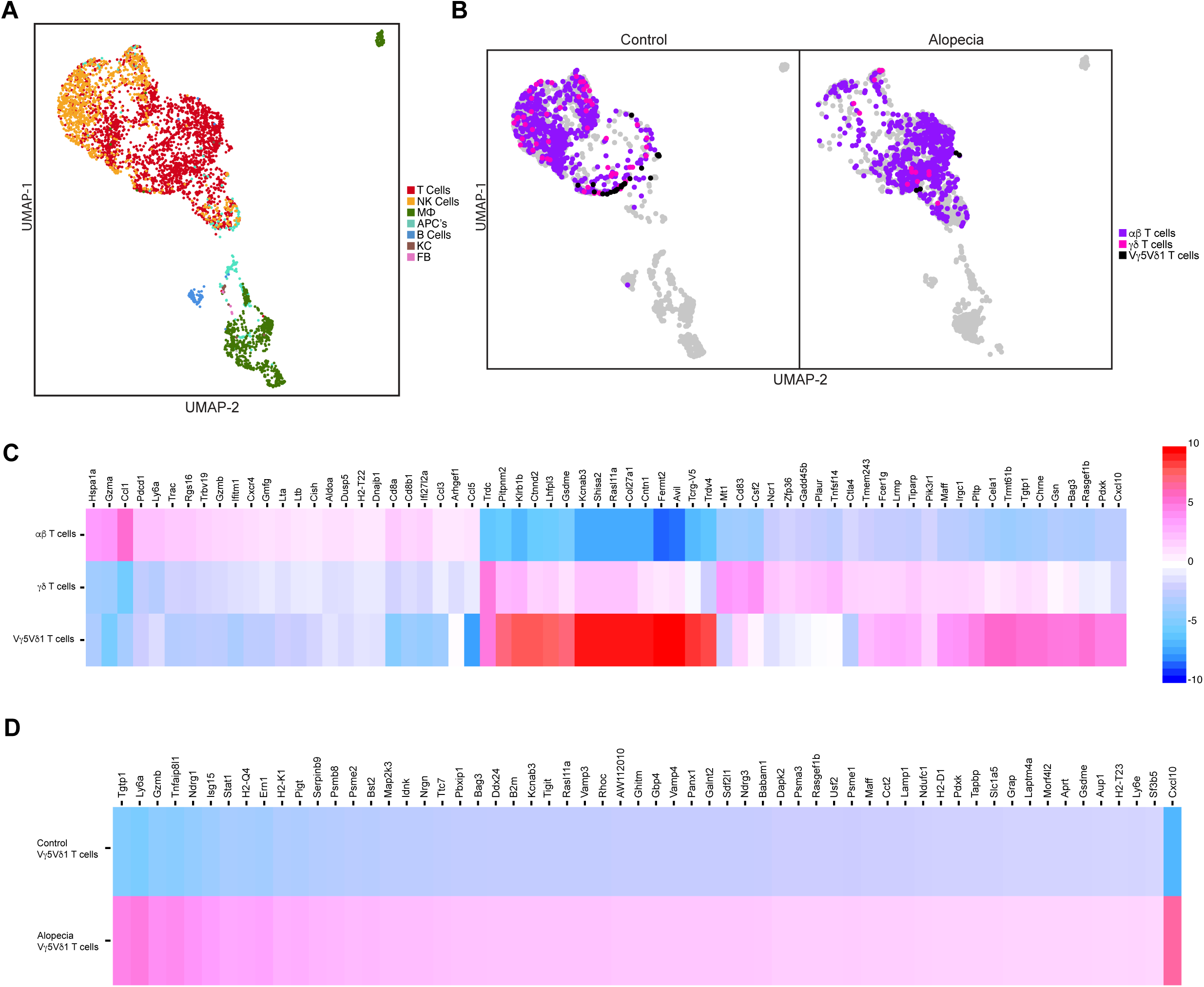
Transcriptional profiles of murine epidermal γδ T cells in AA reveal an increase in genes involved with antiviral activity. Publicly available scRNA sequencing data from Borcherding et al. (a) UMAP plot of the C3H/HeJ murine skin cells from control (n = 2195) and alopecia areata (n = 1946). (b) UMAP plot displaying the relative distribution of αβ T cells, γδ T cells, and Vγ5Vδ1 T cells in control versus alopecia areata. (c) Heatmap rendering of top differentially expressed genes by αβ T cells, γδ T cells, and Vγ5Vδ1 T cell clusters in AA. (d) Heatmap rendering of Vγ5Vδ T cell clusters in control versus AA.

To better understand the roles on skin T cell subsets in disease, we conducted a three-way comparison of upregulated genes from the refined skin T cell subpopulations during alopecia areata to identify differential functions (Fig. 1C). The top genes differentially expressed by skin αβ T cells in alopecia areata were associated with cytotoxic CD8^+^ T cells promoting inflammation (*Hspa1a, Gzma, Ccl1, Ly6a*). This correlates with findings that infiltrating CD8^+^ αβ T cells are heavily implicated in the onset and pathogenesis of alopecia areata (Bertolini et al. 2020). However, it is important to note that some T cell regulatory genes were also expressed uniquely by the αβ T cell population (*Pdcd1, Rgs16*) validating that cytotoxic, inflammatory and regulatory αβ T cells are present during alopecia areata. The top genes differentially expressed by the skin γδ T cell population in alopecia areata were unique from αβ T cells and mostly involved in regulating immune homeostasis (*Mt1, Zfp36, Ctla4*) and proliferation (*Fcer1g, Tnfsf14*). Lastly, genes differentially expressed by epidermis resident Vγ5Vδ1 T cells include T cell recruitment (*Cxcl10)* and vitamin B6 availability (*Pdxk),* implicating these cells in the regulation of inflammatory responses.

To better understand how the function of each cell population changes in alopecia areata, we compared control and alopecia areata samples in a two-way comparison. Not shown is the αβ T cell population, to which the top upregulated genes in alopecia areata are mostly involved in antimicrobial activity (*Ifitm1*, *Ly6a*) and cytolysis (*Gzma*, *Gzmb*), and includes the chemokine signature identified in pathogen-specific effector T cells (*Ccl3*, *Ccl1*, *Ccl4*) (Eberlein et al. 2020). Moreover, the alopecia areata γδ T cell population expresses genes involved in similar biological processes as the αβ T cell population such as antiviral and antimicrobial activity (*Ifitm1*, *Ly6a*, *Ifi27l2a*) and recruitment and activation of monocytes and macrophages (*Cxcl10*, *Ccl3*, *Ccl4*). However, the γδ T cell population also expresses genes involved in the regulation and response of the immune system (*Ctla4*, *Mt1*, *Spp1*) (Fig. 1D).

### Epidermal γδ T cells express antiviral response genes including BST2 in alopecia areata and utilize signaling pathways involved in cell proliferation and migration

Although epidermal γδ T cells were limited in number in the dataset (25 control cells vs. 5 in alopecia areata), we focused on genes expressed in ≥75% of cells from the alopecia areata group (Fig. 2A). These included MHC class I genes (H2-Q4, H2-K1, H2-D1, Tapbpl, B2m) and interferon response genes (Bst2, Ly6a, Tgtp1, Tgtp2, Gzmb, Cxcl10, Stat1). BST2 (also known as tetherin) has not previously been implicated in alopecia areata. Further analysis revealed upregulation of BST2 across T cells and APCs, similar in magnitude to MHC class I genes. To gain insight into functional pathways, we submitted DEGs from epidermal γδ T cells to Ingenuity Pathway Analysis (IPA). Canonical pathways significantly activated included FAK Signaling, CREB Signaling in Neurons, CCR5 Signaling in Macrophages, TEC Kinase Signaling, Phospholipase C Signaling, and PKCθ Signaling in T Lymphocytes (Fig. 2B). These pathways regulate T cell proliferation, survival, and migration. Only one pathway, Inhibition of Matrix Metalloproteases, was downregulated. Overall, these findings suggest that epidermal γδ T cells in alopecia areata may actively proliferate and migrate within the skin.

**Figure 2.**
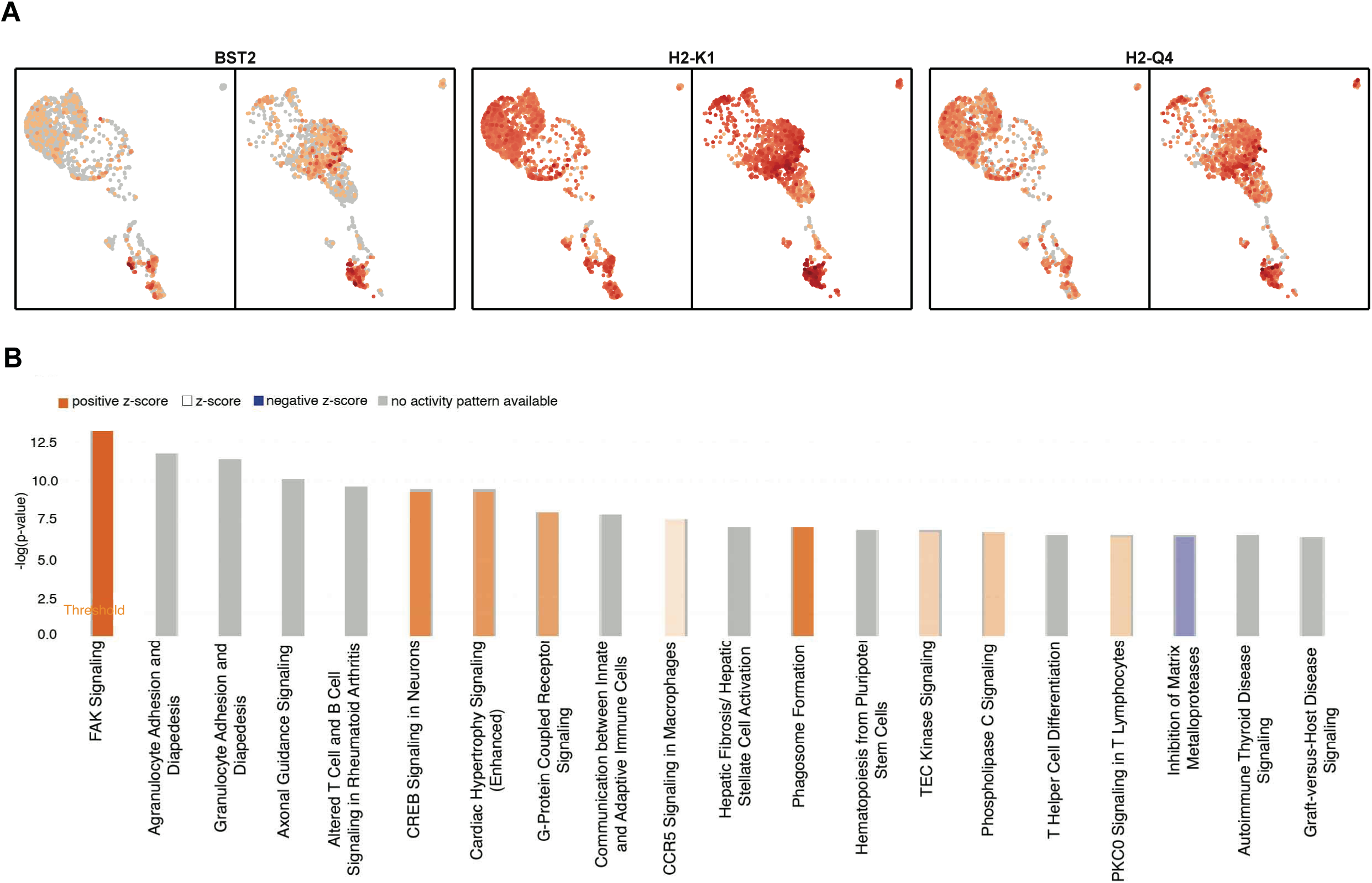
Canonical pathway analysis of all the genes differentially expressed by epidermal γδ T cells comparing alopecia areata to control. (a) Top differentially expressed genes on γδ T cells from publicly available scRNA sequencing data from Borcherding et al. (b) The top differentially expressed genes between alopecia and non-alopecia γδ T cells were clustered into canonical pathways based on IPA knowledge base platform. The threshold (orange line) depicts a default p-value of 0.05. A positive z-score (orange) shows the pathway is activated. A negative z-score (blue) shows the pathway is inhibited. No z-score (white) shows the pathway is neither activated nor inhibited. Z-scores not calculated (grey) shows the pathway did not have sufficient activity for a calculation. Bars are arranged by statistical significance.

### Epidermal γδ T cells and keratinocytes are increased in number in C3H/HeJ mice with alopecia areata

To determine whether epidermal γδ T cells increase or migrate out of the epidermis during alopecia areata, we performed immunofluorescent analysis of C3H/HeJ mouse skin. We quantified γδ T cells in the epidermis of mice with or without alopecia areata. γδ T cells were significantly increased in the epidermis of alopecia areata mice as compared to controls (Fig. 3A, 3B). There were no γδ T cells migrating out of the epidermis to the dermis, so any cellular migration by epidermal γδ T cells in alopecia areata must be occurring within the epidermal compartment. γδ T cells play key roles in epidermal homeostasis and wound repair through the production of growth factors so we quantified the number of keratinocytes (Sharp et al. 2005).

**Figure 3.**
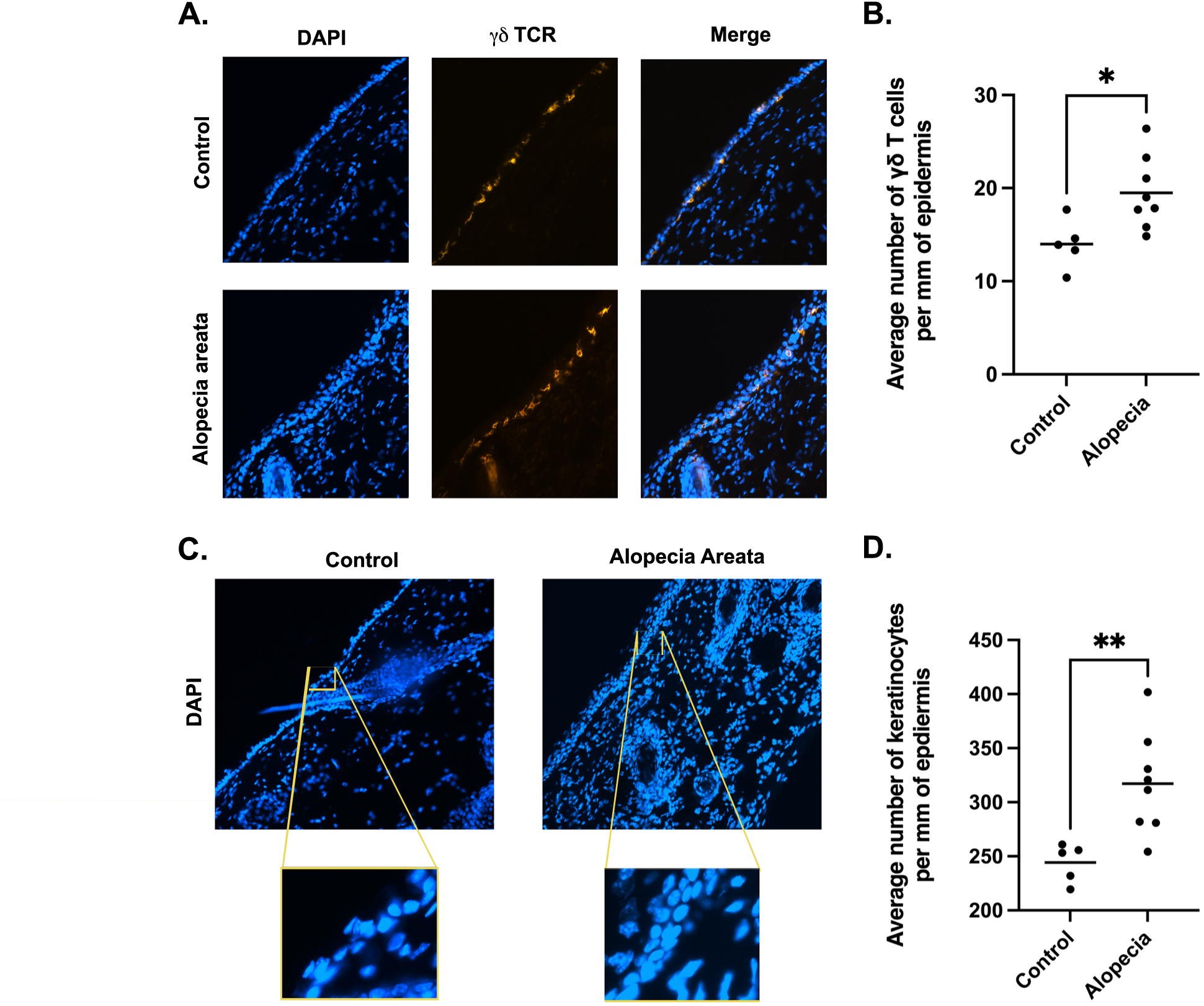
Epidermal γδ T cells and keratinocytes are elevated in the skin of mice with alopecia areata. (a) Immunofluorescence images of the average number of epidermal γδ T cells from skin lesions in C3H/HeJ mice with (n=8) and without (n=5) alopecia areata. (b) Quantitation of epidermal γδ T cells in sections with and without alopecia areata. (c) Immunofluorescent images of the average number of keratinocytes from C3H/HeJ mice with (n=8) and without (n=5) alopecia areata. (d) Quantitation of keratinocytes in sections with and without alopecia areata. Significance determined using an unpaired t-test (*p < .05,**p < .01).

We found a significant increase in keratinocytes in the epidermis of mice with alopecia areata as compared to controls (Fig. 3C, 3D). Our findings support a model in which signaling through the FAK, TEC Kinase, G-Protein Receptor and phospholipase C signaling pathways promotes coordinated proliferation of both epidermal γδ T cells and keratinocytes in alopecia areata.

### BST2 is upregulated on epidermal γδ T cells near hair follicles in mice with alopecia areata

To validate the scRNAseq data identifying *BST2* expression by epidermal γδ T cells, we examined skin from mice with alopecia areata to determine whether BST2 is upregulated by epidermal γδ T cells. Our analysis revealed that epidermal γδ T cells express BST2 in alopecia areata, particularly near hair follicles (Fig. 4A). To better understand the function of BST2 in epidermal γδ T cells we further analyzed BST2^+^ and BST2^−^ epidermal γδ T cells in the scRNA-seq data to identify differentially expressed genes (Fig. 4B). The results further validate that ΒST2^+^ epidermal γδ T cells are involved in antiviral responses and interferon signaling (*Isg15*, *Stat1)*. In contrast, BST2^-^ cells express genes involved in immune regulation and moderation (*CD74, Pdcd1*, *Nfkbid, Il7r*), as well as tissue repair and regeneration (*Areg*).

**Figure 4.**
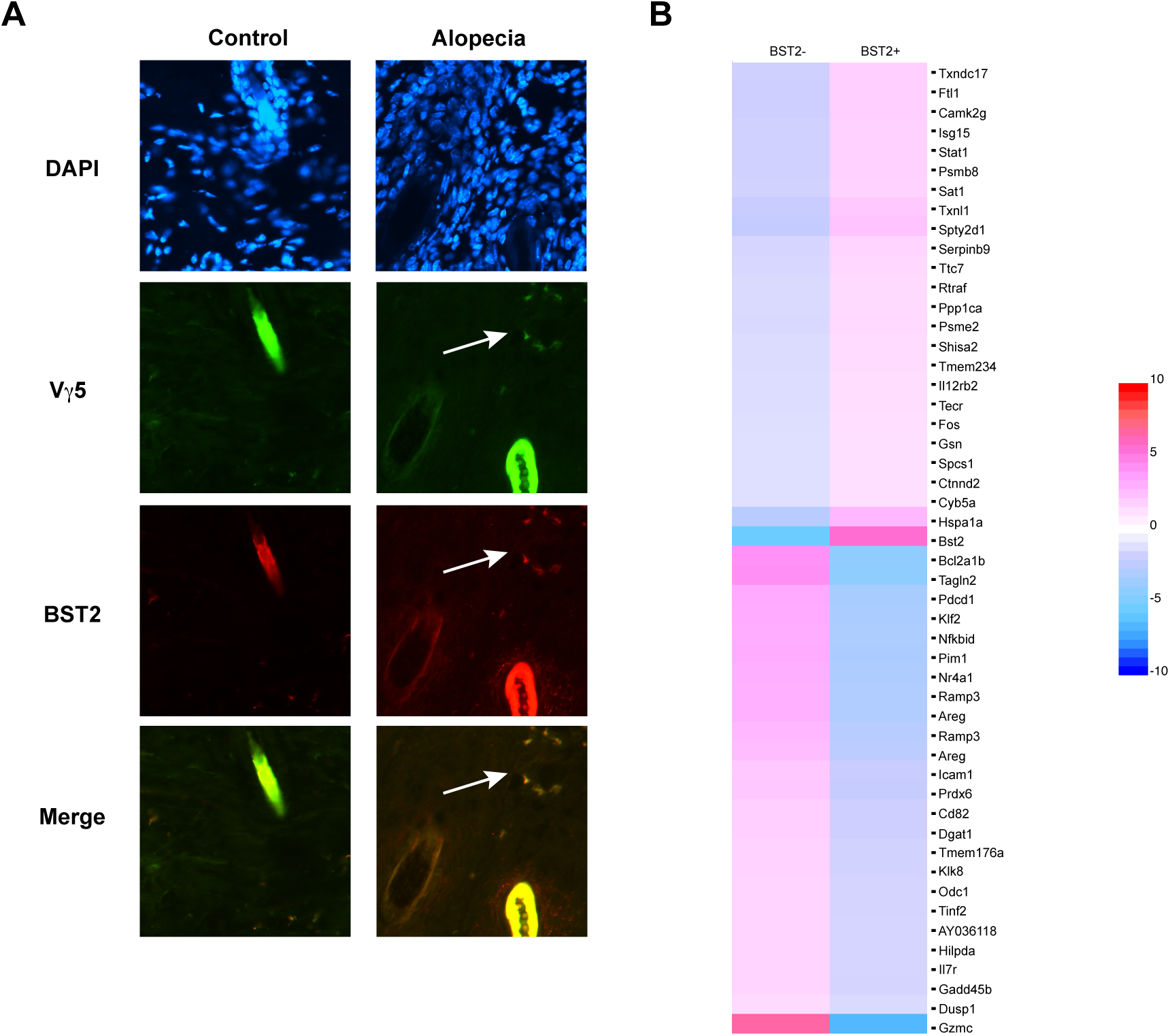
BST2 expression is increased on epidermal γδ T cells and primarily localized near hair follicles in mice with alopecia areata. (a) Representative immunofluorescent images of cross-sections from C3H/HeJ mice show epidermal γδ T cells surrounding hair follicles in alopecia areata (n=3). (b) Heatmap rendering of BST2^+^ and BST2^-^ epidermal γδ T cell populations.

### BST2 distinguishes IFN-γ production by CD8 T cells in alopecia areata

The scRNAseq dataset also revealed an increase in CD8 cells (*Cd3e*, *Cd3d*, *Cd3g*, *Cd8a*/*Cd8b1*, with no *Cd4* expression) expressing *BST2*. We quantified BST2^+^ CD8 T cells in the skin of both alopecia areata and control mice using immunofluorescent microscopy (Fig. 5A). While there were very few detectable BST2^+^ CD8 T cells in control mouse skin, the number of BST2^+^ CD8 T cells was increased in mice with alopecia areata (Fig. 5B). To determine whether BST2 expressing CD8 T cells exhibit similar gene expression as epidermal γδ T cells expressing BST2, we analyzed CD8 T cells with and without BST2 expression in the scRNA-seq dataset to infer the function for these cell subsets in alopecia areata (Fig. 5C). BST2^+^ CD8 T cells exhibit high expression of genes associated with antiviral responses and IFN-γ signaling (*Ifng, Xcl1, Ly6a, Ly6e, Ifi27l2a)*. Unique to CD8 T cells, BST2^+^ CD8 T cells express *IL-13*, which has been identified in genome wide association studies of alopecia areata patients (Cai et al. 2011).

**Figure 5.**
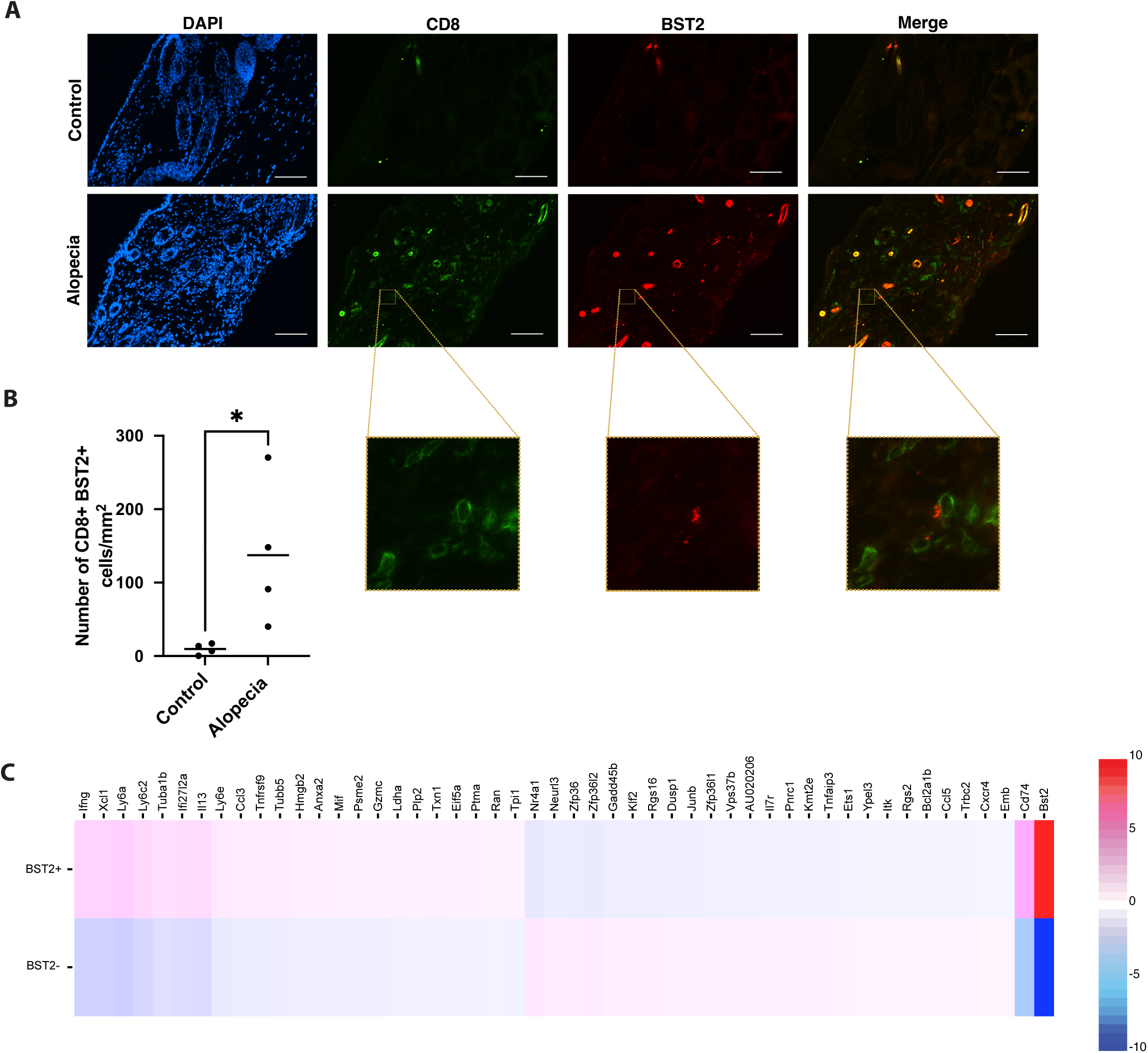
BST2 expression by CD8 T cells is elevated in the skin of mice with alopecia areata. (a) Immunofluorescent images were used to quantify the number of CD8^+^BST2^+^ cells in the dermis from skin samples of C3H/HeJ mice with (n=4) and without (n=4) alopecia areata. (b) Quantification of CD8+ cells in the dermis of C3H/HeJ mice with and without alopecia areata. (c) Heatmap of top differentially expressed genes by BST2^+^CD8^+^ versus BST2^-^CD8^+^ T cells. Significance determined using an unpaired t-test (*p < 0.05).

### BST2-expressing CD11b^+^ macrophages are increased in the skin during alopecia areata and express antiviral response and complement system genes

We also observed upregulation of BST2 on macrophages (Cd14^+^, Lyz2^+^), predominantly expressing CD11b (Itgam) (Fig. 1A, Suppl. Fig. 1). Immunofluorescence revealed an elevated number of dermal CD11b^+^ cells in alopecia areata (p = 0.07), with a significant increase in BST2^+^ CD11b^+^ macrophages (Fig. 6A–D). Differential gene expression analysis showed that BST2^+^ macrophages upregulate antiviral and interferon response genes (Ccl8, Ifitm1, Cxcl3, Ifit2) (Suppl. Table 1). Ingenuity Pathway Analysis of BST2^+^ macrophages identified enrichment of pathways related to pathogen sensing, cytokine storm signaling, and MAPK signaling in influenza (Fig. 6E). Additionally, these cells express complement genes (C1qa, C1qb, C1qc) and M2 macrophage-associated transcripts (Apoe, Tmem176b, Hp), indicating an anti-inflammatory or reparative macrophage phenotype potentially driven by IL-17 signaling.

**Figure 6.**
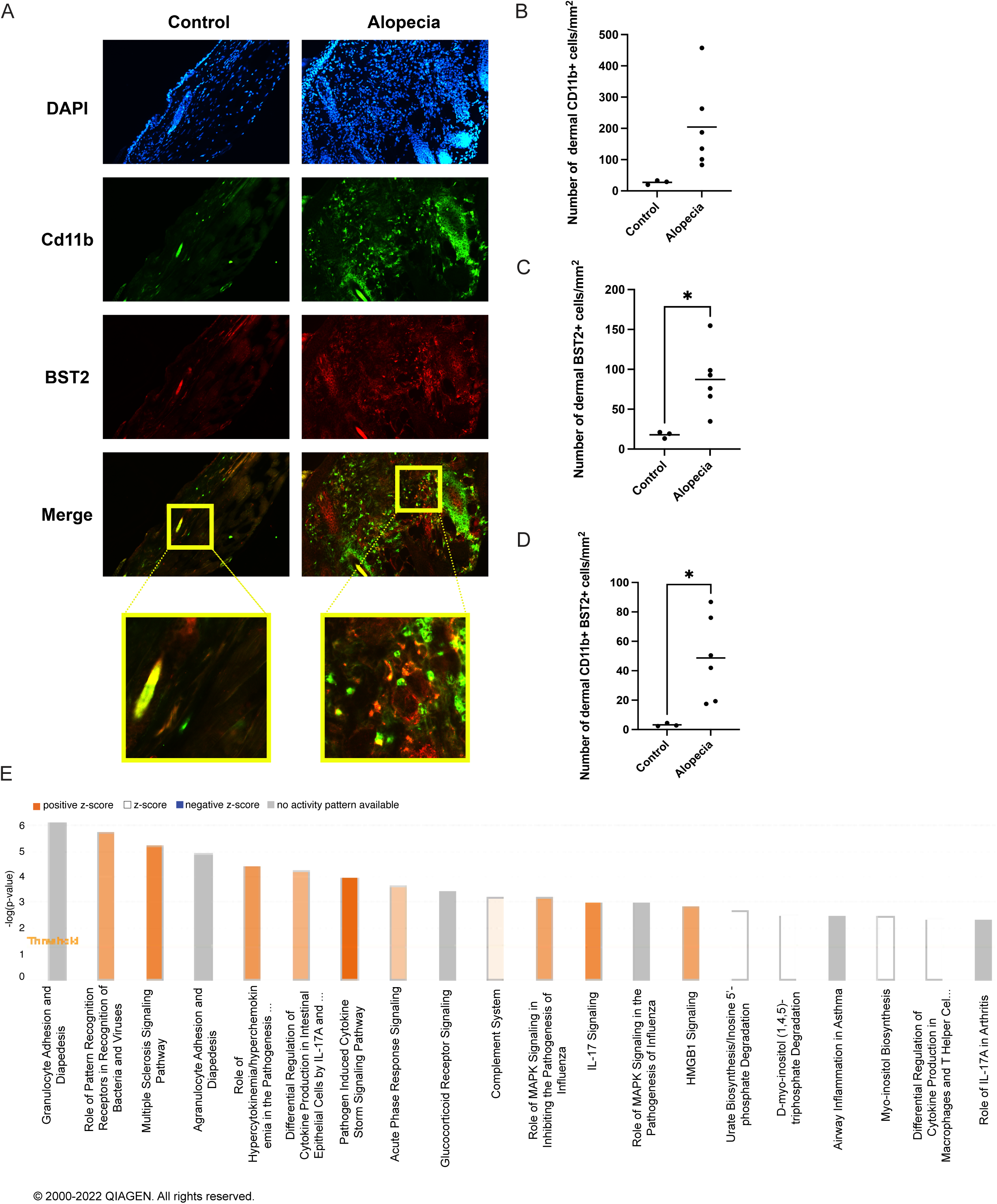
BST2-expressing CD11b^+^ macrophages are increased in the skin during alopecia areata and express genes involved in antiviral responses and the complement system. (a) Immunofluorescent images were used to quantify the number of CD11b^+^, BST2^+^ and CD11b^+^ BST2^+^ cells from skin samples of C3H/HeJ mice with (n=6) and without (n=3) alopecia areata. (b) Quantification of CD11b^+^ cells. (c) Quantification of BST2^+^ cells. (d) Quantification of CD11b^+^BST2^+^ cells. (e) Canonical pathway analysis of the top differentially expressed genes using IPA sorted by BST2^+^ and BST2^-^ Cd11b^+^ cells. Significance determined using an unpaired t-test (*p < 0.05).

### BST2 is expressed by patients with alopecia areata and downregulated with Tofacitinib

To assess translational reference of BST2, we reanalyzed publicly available bulk RNAseq data from alopecia areata patients before and after treatment with the JAK inhibitor tofacitinib (Crispin et. al, 2016; (Gay et al. 2013). Genes downregulated post-treatment include *Bst2*, *Ccl27*, *Ifi6*, *Ifi27*, and *Ccl13*, while keratin-associated genes are upregulated (Table 1). These findings are consistent with mouse data and suggest that BST2 may serve as a disease biomarker that is responsive to immunomodulatory therapy.

**Table 1.** Reanalyzed RNA-seq data identifies differentially expressed genes, including BST2, post-tofacitinib treatment (Crispen et al., 2016).

| Gene | Log2FC | p-value | Gene | Log2FC | p-value |
| --- | --- | --- | --- | --- | --- |
| *Upregulated* |  |  | *Downregulated* |  |  |
| *KRTAP5-2* | 0.798 | 6.76E-07 | *HBA1/HBA2* | -0.920 | 2.95E-10 |
| *KRTAP8-1* | 0.773 | 6.59E-07 | *CCL27* | -0.695 | 1.20E-10 |
| *KRTAP1-3* | 0.736 | 2.12E-06 | *IFI6* | -0.646 | 3.73E-09 |
| *LYG2* | 0.732 | 5.89E-06 | *ODF3B* | -0.578 | 1.11E-07 |
| *KRTAP19-3* | 0.773 | 6.55E-06 | *BST2* | -0.549 | 2.12E-06 |
| *KRT86* | 0.718 | 3.51E-06 | *LGALS7/LGALS7* | -0.548 | 6.65E-04 |
| *KRTAP4-5* | 0.704 | 1.26E-05 | *NDUFA13* | -0.527 | 1.16E-04 |
| *KRTAP12-2* | 0.694 | 1.27E-05 | *IFI27* | -0.525 | 1.96E-05 |
| *KRT82* | 0.677 | 2.08E-05 | *CCL13* | -0.508 | 1.86E-04 |
| *KRT33A* | 0.667 | 2.32E-05 | *TYMP* | -0.505 | 5.85E-04 |

## Discussion

Infiltrating CD8 cytotoxic T cells, CD4 helper T cells, and NK cells are known to be key mediators in the pathogenesis of alopecia areata. In this study, we further validate that αβ T cells in the C3H/HeJ mouse model of alopecia areata exhibit genes indicative of cytolytic, pathogen-specific effector cells such as Gzma, Gzmb, CCL1, and CCL3. Similarly, CD8 effectors producing CCL1, CCL3 and CCL4 have been identified in viral infections such as LCMV and VSV (Eberlein et al. 2020). Further, CD8 cytotoxic T cell “swarming” around tumor cells is driven by CCL3 and CCL4, a phenomenon reminiscent of the “swarm of bees” infiltration pattern around hair follicles in alopecia areata (Zhao and Guan 2011). Recent studies have begun to highlight the role of tissue-resident T cell populations, such as γδ T cells, may recruit effector populations such as CD8 T cells in alopecia areata. In humans, Vδ1 T cells have been identified near the hair follicle infundibulum with elevated numbers inside the proximal epithelium in nonlesional and lesional skin from patients with alopecia areata (Jabbari et al. 2016). Here we also find elevated numbers of epidermal γδ T cells in the C3H/HeJ mouse model of alopecia areata. These epidermal γδ T cells alone or grouped with dermal γδ T cells express CXCL10, which has been linked to the early infiltration of leukocytes into the skin during the development of alopecia areata (Boismenu and Havran 1994).

Epidermal γδ T cells originate in the fetal thymus and seed the epidermis, where they maintain long-term residency through self-renewal (Havran and Allison 1990; Jameson et al. 2004). In alopecia areata, this homeostatic balance appears disrupted, resulting in elevated numbers of epidermal γδ T cells. Gene pathway analysis of these cells revealed enrichment in signaling cascades associated with proliferation, particularly focal adhesion kinase (FAK) signaling. FAK is a downstream effector of T cell receptor engagement and plays critical roles in cell cycle and proliferation (Chapman and Houtman 2014; Zhao et al. 1998). Further analysis of the top differentially expressed genes of epidermal γδ T cells in alopecia areata reveal BST2, an antiviral factor, as a biomarker of interest. BST2 is a transmembrane protein that functions by tethering viral particles to the cell surface (Hammonds et al. 2010). BST2^+^ epidermal γδ T cells are uniquely associated with the hair follicle suggesting that epidermal γδ T cells may play roles in identifying infection at the hair follicle. Furthermore, the top DEGs expressed by epidermal γδ T cells are heavily correlated with anti-viral activity and MHC class I presentation. Emerging research suggests alopecia areata can develop during or as a result of viral infections such as with swine flu, hepatitis virus (hepatitis B and hepatitis C), and more recently, SARS-CoV-2 (Alotaibi et al. 2023; Christensen and Jafferany 2022; Ito and Tokura 2012; Wiwanitkit and Wiwanitkit 2014).

In alopecia areata, the hair follicle immune privilege collapses after attack from infiltrating CD8^+^ NKG2D^+^ T cells and natural killer cells (Bertolini et al. 2020). In addition to γδ T cells, we found increased expression of BST2 in CD8^+^ T cells localized around the hair follicles in alopecia areata. This suggests a potential amplification loop whereby early interferon responses from tissue-resident cells may prime cytotoxic T cells to accumulate at the hair follicle. Transcriptional analysis of BST2^+^ CD8^+^ T cells revealed upregulation of *IFNG* and *Xcl1*. IFN-γ is a critical antiviral cytokine and a key activator of the JAK/STAT pathway (Horvath 2004; Tau and Rothman 1999). Dermal macrophages also contributed to the interferon-rich microenvironment, with a significant increase in CD11b^+^BST2^+^ macrophages in alopecia areata skin. These cells expressed antiviral and interferon response genes and were enriched for pathways involved in innate immune activation, complement signaling, and cytokine response.

Interestingly, prior studies show that depleting macrophages in the skin can promote hair follicle regeneration and enhance hair growth (Castellana et al. 2014). This suggests that while macrophages contribute to immune defense, they may also perpetuate inflammatory signaling that disrupts follicular homeostasis.

Macrophages and T cells together play important roles in regulating hair follicle cycling and regeneration. Macrophages secrete cytokines and growth factors to influence hair cycle transitions (Wang et al. 2019). Conversely, macrophages can inhibit hair follicle growth via the JAK-signal transducer and activator of transcription (STAT) pathway (Triyangkulsri and Suchonwanit 2018). JAK inhibitors are an FDA-approved treatment for alopecia areata. In our analysis of publicly available data from the skin of human patients with alopecia areata, we observe elevated BST2 expression that is reduced following treatment with the JAK inhibitor, tofacitinib (Crispin et al. 2016). This suggests that BST2 expression may serve as a functional readout of IFN-γ–driven JAK/STAT activation in alopecia areata.

Taken together, this data highlights a potential model in which epidermal γδ T cells initiate an antiviral response, upregulating BST2 and chemokines such as CXCL10, thereby recruiting CD8^+^ T cells and macrophages to the hair follicle. These infiltrating cells further amplify interferon signaling, promote immune privilege collapse, and drive pathologic hair loss.

Targeting the pathways regulating BST2 expression and IFN-γ–JAK/STAT signaling may offer novel therapeutic strategies for restoring immune tolerance and follicular homeostasis in alopecia areata.

## Materials and Methods

### Mice

Female *C3H/HeJ* mice were purchased from The Jackson Laboratory (Bar Harbor, ME) and maintained under specific pathogen-free conditions in the animal facility at California State University, San Marcos. All mice were between 6 to 24 months of age and were provided with food and water *ad libitum*. All animal procedures involving animals were reviewed and approved by the Institutional Animal Care and Use Committee of California State University San Marcos (protocol 22-001).

### Single cell RNA sequencing data processing and analysis

Publicly available scRNAseq data from mouse skin with and without alopecia areata were obtained from the National Center for Biotechnology Information (NCBI) Gene Expression Omnibus (GEO) under GEO accession numbers GSE145095 and GSE68801 (Borcherding et al. 2022). The raw data was demultiplexed and aligned to the mouse reference genome (mm10) using the Cell Ranger (v5.0.1) pipeline (10X Genomics). The study by Borcherding isolated skin infiltrating CD45^+^ cells from C3H/HeJ mice with and without alopecia areata (Borcherding et al. 2022). Cellranger was used to generate a cloupe file that was uploaded to Loupe Browser v6.0 (10x Genomics) for downstream analysis. After normalization, 37,574 mean reads per cell and 1,345 median genes per cell were detected from a total of 4,141 cells. Uniform manifold approximation projections (UMAP) were used to visualize spatial gene expression data and differentially expressed genes (DEGs) were calculated using the Locally Distinguishing tool in Loupe. T cells were clustered into primary subsets (αβ T cells, γδ T cells, and Vγ5Vδ1 T cells). Vγ5Vδ1 T cell gene expression between alopecia and control samples was evaluated and assessed by Ingenuity Pathway Analysis (IPA) (Qiagen) (Venlo, Netherlands). BST2 expressing populations were compared to non-expressing populations in Vγ5 T cells, CD8 T cells, and CD11b^+^ macrophages and at times evaluated using IPA (Qiagen).

### Tissue preparation and immunofluorescent microscopy

Skin was acquired from C3H/HeJ mice with alopecia areata from regions lacking hair. Control skin samples were obtained from age-matched C3H/HeJ mice without alopecia areata. Skin sections measuring 2mm x 2mm were placed midline down in cryomolds, frozen using Optimal Cutting Temperature compound on dry ice, and stored at -80 C. Embedded skin was sectioned using a cryostat (Leica CM 1520, Deer Park, IL). Skin sections were cut at 10 μm thickness, fixed for 10 minutes with 4% paraformaldehyde in PBS, and blocked for 30 minutes using gelatin block solution (2.5% Normal Goat Serum, 2.5% Normal Donkey Serum, 1% BSA, 2% fish gelatin, 0.1% Triton X, 0.75% glycine). Antibodies were purchased from either BioLegend (San Diego, CA) or BD Biosciences (Franklin Lakes, NJ). Sections were stained with anti-CD3ε (10 ug/mL, clone 145-2C11), anti-Vγ3 TCR (2 ug/mL, 536) anti-CD8 (4 ug/mL, YTS156.7.7), anti-CD317 (8 ug/mL, 129c), anti-CD11b (5 ug/mL, 5C6) or anti-γδTCR (4ug/mL, GL3) for 1 hour, washed in 1x PBS, and mounted using SlowFade Gold Antifade Mounting Media with DAPI (Thermo Fisher, San Diego, CA). A minimum of 4 sections per mouse and 25 digital images per section were acquired for each mouse using an immunofluorescent microscope (Nikon DS-Qi2, Nikon, Tokyo, JPN) at x200 magnification. Images were captured using NIS Element D 4.1300 64 program (Nikon) and analyzed in Adobe® Photoshop® software (Adobe, San Jose, CA). Epidermis length and cell number were quantified using ImageJ software (National Institutes of Health, Bethesda, MD). Dermal cell number was quantified using mm^2^ grids using Adobe® Photoshop® software (Adobe).

### Analysis of publicly available patient bulk RNA seq data

Publicly available RNAseq data collected from skin biopsies of alopecia areata patients prior to and post treatment with tofacitinib citrate were retrieved from GEO accession number GSE80688 in the NCBI Gene Expression Omnibus (Crispin et al. 2016). Differential gene expression analysis was performed using DEseq2 within the Galaxy platform, comparing baseline and 8 week post-tofacitinib-citrate treatment samples. Additionally, data was compared to a healthy human gene sequence also obtained from NCBI. Genes were ranked based on fold change (log2(Fold Change)) and the top ten upregulated and downregulated genes were identified using IPA (Qiagen).

### Statistical analysis

Statistical analysis was performed using GraphPad Prism (version 10; Dotmatics, Boston, MA). A parametric, unpaired, Student’s t-test was used to determine significance for average number of keratinocytes, γδ T cells, CD8^+^ T cells, and CD11b^+^ macrophages. Log2-fold change was calculated in Loupe Browser by using the ratio of normalized mean gene unique molecular identifiers (UMI) counts in each cluster relative to all other clusters. IPA uses Ingenuity Knowledge Base to calculate log 2-fold change and the statistical significance.

## Ethics Statement

Collection of animal tissue samples for this study was approved as part of the study protocol. This animal study was approved by CSUSM IACUC - approval: 22-001.

## Data Availability Statement

Murine transcriptomic data from control and alopecia areata skin were obtained from Borcherding et al. through the NCBI Gene Expression Omnibus (https://www.ncbi.nlm.nih.gov/geo/query/acc.cgi) under acquisition numbers GSE145095 and GSE68801. Additionally, bulk RNA-sequencing data of alopecia areata patients before and after treatment with tofacitinib as reported by Crispin et al. was obtained from GEO under accession number GSE80688.

## Acknowledgements

The authors thank the Jameson lab members for manuscript review.

## Credit Statement

Conceptualization: AG, MM, JJ; Formal Analysis: JC, AG, CV, MK, YV, MM, AS, AK, JJ; Methodology: JC, AG, CV, MK, YV, MM, AS, AK, JJ; Data curation: JC, AG, CV, MK, YV, MM, AK; Writing - Original Draft Presentation: JC, AG, JJ; Writing - Review and Editing: JC, AG, CV, MK, YV, MM, AS, AK, JJ

## References

Alotaibi MA, Altaymani A, Al-Omair A, Alghamdi W. Viral-Induced Rapidly Progressive Alopecia Universalis: A Case Report and Literature Review. Cureus. 2023;15(4):e37406

Bertolini M, McElwee K, Gilhar A, Bulfone-Paus S, Paus R. Hair follicle immune privilege and its collapse in alopecia areata. Exp Dermatol. 2020;29(8):703–25

Boismenu R, Havran WL. Modulation of epithelial cell growth by intraepithelial gamma delta T cells. Science. 1994;266(5188):1253–5

Borcherding N, Crotts SB, Ortolan LS, Henderson N, Bormann NL, Jabbari A. A transcriptomic map of murine and human alopecia areata. JCI Insight. 2022;5(13):e137424

Cai Y, Shen X, Ding C, Qi C, Li K, Li X, et al. Pivotal role of dermal IL-17-producing γδ T cells in skin inflammation. Immunity. 2011;35(4):596–610

Castellana D, Paus R, Perez-Moreno M. Macrophages Contribute to the Cyclic Activation of Adult Hair Follicle Stem Cells. PLoS Biol. 2014;12(12):e1002002

Chapman NM, Houtman JCD. Functions of the FAK family kinases in T cells: Beyond actin cytoskeletal rearrangement. Immunol Res. 2014;59(0):23–34

Christensen RE, Jafferany M. Association between alopecia areata and COVID-19: A systematic review. JAAD Int. 2022;7:57–61

Connell SJ, Jabbari A. The current state of knowledge of the immune ecosystem in alopecia areata. Autoimmun Rev. 2022;21(5):103061

Crispin MK, Ko JM, Craiglow BG, Li S, Shankar G, Urban JR, et al. Safety and efficacy of the JAK inhibitor tofacitinib citrate in patients with alopecia areata. JCI Insight. American Society for Clinical Investigation; 2016;1(15) Available from: https://insight.jci.org/articles/view/89776

Eberlein J, Davenport B, Nguyen TT, Victorino F, Jhun K, van der Heide V, et al. Chemokine Signatures Of Pathogen-Specific T Cells I: Effector T Cells. J Immunol. 2020;205(8):2169–87

Gay D, Kwon O, Zhang Z, Spata M, Plikus MV, Holler PD, et al. Fgf9 from dermal γδ T cells induces hair follicle neogenesis after wounding. Nat Med. Nature Publishing Group; 2013;19(7):916–23

Gilhar A, Ullmann Y, Berkutzki T, Assy B, Kalish RS. Autoimmune hair loss (alopecia areata) transferred by T lymphocytes to human scalp explants on SCID mice. J Clin Invest. 1998;101(1):62–7

Hammonds J, Wang J-J, Yi H, Spearman P. Immunoelectron Microscopic Evidence for Tetherin/BST2 as the Physical Bridge between HIV-1 Virions and the Plasma Membrane. PLoS Pathog. 2010;6(2):e1000749

Han Y-M, Sheng Y-Y, Xu F, Qi S-S, Liu X-J, Hu R-M, et al. Imbalance of T-helper 17 and regulatory T cells in patients with alopecia areata. The Journal of Dermatology. 2015;42(10):981–8

Havran WL, Allison JP. Origin of Thy-1+ dendritic epidermal cells of adult mice from fetal thymic precursors. Nature. 1990;344(6261):68–70

Horvath CM. The Jak-STAT Pathway Stimulated by Interferon γ. Sci. STKE. 2004;2004(260) Available from: https://www.science.org/doi/10.1126/stke.2602004tr8

Ito T, Tokura Y. Alopecia areata triggered or exacerbated by swine flu virus infection. The Journal of Dermatology. 2012;39(10):863–4

Jabbari A, Nguyen N, Cerise JE, Ulerio G, de Jong A, Clynes R, et al. Treatment of an alopecia areata patient with tofacitinib results in regrowth of hair and changes in serum and skin biomarkers. Exp Dermatol. 2016;25(8):642–3

Jagielska D, Redler S, Brockschmidt FF, Herold C, Pasternack SM, Garcia Bartels N, et al. Follow-Up Study of the First Genome-Wide Association Scan in Alopecia Areata: IL13 and KIAA0350 as Susceptibility Loci Supported with Genome-Wide Significance. Journal of Investigative Dermatology. 2012;132(9):2192–7

Jameson JM, Cauvi G, Witherden DA, Havran WL. A keratinocyte-responsive gamma delta TCR is necessary for dendritic epidermal T cell activation by damaged keratinocytes and maintenance in the epidermis. J Immunol. 2004;172(6):3573–9

John KK-G, Brockschmidt FF, Redler S, Herold C, Hanneken S, Eigelshoven S, et al. Genetic Variants in CTLA4 Are Strongly Associated with Alopecia Areata. Journal of Investigative Dermatology. 2011;131(5):1169–72

Johnson MD, Witherden DA, Havran WL. The Role of Tissue-resident γδ T Cells in Stress Surveillance and Tissue Maintenance. Cells. Multidisciplinary Digital Publishing Institute; 2020;9(3):686

Kwon O, Senna MM, Sinclair R, Ito T, Dutronc Y, Lin C-Y, et al. Efficacy and Safety of Baricitinib in Patients with Severe Alopecia Areata over 52 Weeks of Continuous Therapy in Two Phase III Trials (BRAVE-AA1 and BRAVE-AA2). Am J Clin Dermatol. 2023;1–9

Lee HH, Gwillim E, Patel KR, Hua T, Rastogi S, Ibler E, et al. Epidemiology of alopecia areata, ophiasis, totalis, and universalis: A systematic review and meta-analysis. Journal of the American Academy of Dermatology. Elsevier; 2020;82(3):675–82

Martinez-Mir A, Zlotogorski A, Gordon D, Petukhova L, Mo J, Gilliam TC, et al. Genome-wide Scan for Linkage Reveals Evidence of Several Susceptibility Loci for Alopecia Areata. Am J Hum Genet. 2007;80(2):316–28

McElwee KJ, Freyschmidt-Paul P, Hoffmann R, Kissling S, Hummel S, Vitacolonna M, et al. Transfer of CD8+ Cells Induces Localized Hair Loss Whereas CD4+/CD25− Cells Promote Systemic Alopecia Areata and CD4+/CD25+ Cells Blockade Disease Onset in the C3H/HeJ Mouse Model. Journal of Investigative Dermatology. 2005;124(5):947–57

Nasiri S, Salehi A, Rakhshan A. Infiltration of Mast Cells in Scalp Biopsies of Patients with Alopcia Areata or Androgenic Alopecia Versus Healthy Individuals: A Case Control Study. Galen Med J. 2020;9:e1962

Nielsen MM, Witherden DA, Havran WL. γδ T cells in homeostasis and host defence of epithelial barrier tissues. Nat Rev Immunol. 2017;17(12):733–45

Petukhova L, Duvic M, Hordinsky M, Norris D, Price V, Shimomura Y, et al. Genome-wide association study in alopecia areata implicates both innate and adaptive immunity. Nature. 2010;466(7302):113–7

Ranki A, Kianto U, Kanerva L, Tolvanen E, Johansson E. Immunohistochemical and electron microscopic characterization of the cellular infiltrate in alopecia (areata, totalis, and universalis). J Invest Dermatol. 1984;83(1):7–11

Sharp LL, Jameson JM, Cauvi G, Havran WL. Dendritic epidermal T cells regulate skin homeostasis through local production of insulin-like growth factor 1. Nat Immunol. 2005;6(1):73–9

Simakou T, Butcher JP, Reid S, Henriquez FL. Alopecia areata: A multifactorial autoimmune condition. J Autoimmun. 2019;98:74–85

Tau G, Rothman P. Biologic functions of the IFN-gamma receptors. Allergy. 1999;54(12):1233– 51

Tiwari R, de la Torre JC, McGavern DB, Nayak D. Beyond Tethering the Viral Particles: Immunomodulatory Functions of Tetherin (BST-2). DNA Cell Biol. 2019;38(11):1170–7

Toulon A, Breton L, Taylor KR, Tenenhaus M, Bhavsar D, Lanigan C, et al. A role for human skin–resident T cells in wound healing. J Exp Med. 2009;206(4):743–50

Triyangkulsri K, Suchonwanit P. Role of janus kinase inhibitors in the treatment of alopecia areata. Drug Des Devel Ther. 2018;12:2323–35

Uchida Y, Gherardini J, Schulte-Mecklenbeck A, Alam M, Chéret J, Rossi A, et al. Pro-inflammatory Vδ1+T-cells infiltrates are present in and around the hair bulbs of non-lesional and lesional alopecia areata hair follicles. J Dermatol Sci. 2020;100(2):129–38

Wang ECE, Dai Z, Ferrante AW, Drake CG, Christiano AM. A Subset of TREM2+ Dermal Macrophages Secretes Oncostatin M to Maintain Hair Follicle Stem Cell Quiescence and Inhibit Hair Growth. Cell Stem Cell. 2019;24(4):654–669.e6

Wiwanitkit S, Wiwanitkit V. Alopecia Due to Hepatitis Virus Infections (Hepatitis B and Hepatitis C). tdd. 2014;8(2):101–3

Xing L, Dai Z, Jabbari A, Cerise JE, Higgins CA, Gong W, et al. Alopecia areata is driven by cytotoxic T lymphocytes and is reversed by JAK inhibition. Nat Med. 2014;20(9):1043–9

Zhang B, Zhao Y, Cai Z, Caulloo S, McElwee KJ, Li Y, et al. Early stage alopecia areata is associated with inflammation in the upper dermis and damage to the hair follicle infundibulum: Early alopecia areata pathology. Australasian Journal of Dermatology. 2013;54(3):184–91

Zhao X, Guan J-L. Focal adhesion kinase and its signaling pathways in cell migration and angiogenesis. Adv Drug Deliv Rev. 2011;63(8):610–5

Zhao J-H, Reiske H, Guan J-L. Regulation of the Cell Cycle by Focal Adhesion Kinase. The Journal of Cell Biology. The Rockefeller University Press; 1998;143(7):1997

